# NOGEA: Network-Oriented Gene Entropy Approach for Dissecting Disease Comorbidity and Drug Repositioning

**DOI:** 10.1101/2020.04.01.019901

**Authors:** Zihu Guo, Yingxue Fu, Chao Huang, Chunli Zheng, Ziyin Wu, Xuetong Chen, Shuo Gao, Yaohua Ma, Mohamed Shahen, Yan Li, Pengfei Tu, Jingbo Zhu, Zhenzhong Wang, Wei Xiao, Yonghua Wang

## Abstract

Rapid development of high-throughput technologies has permitted the identification of an increasing number of disease-associated genes (DAGs), which are important for understanding disease initiation and developing precision therapeutics. However, DAGs often contain large amounts of redundant or false positive information, leading to difficulties in quantifying and prioritizing potential relationships between these DAGs and human diseases. In this study, a network-oriented gene entropy approach (NOGEA) is proposed for accurately inferring master genes that contribute to specific diseases by quantitatively calculating their perturbation abilities on directed disease-specific gene networks. In addition, we confirmed that the master genes identified by NOGEA have a high reliability for predicting disease-specific initiation events and progression risk. Master genes may also be used to extract the underlying information of different diseases, thus revealing mechanisms of disease comorbidity. More importantly, approved therapeutic targets are topologically localized in a small neighborhood of master genes on the interactome network, which provides a new way for predicting new drug-disease associations. Through this method, 11 old drugs were newly identified and predicted to be effective for treating pancreatic cancer and then validated by *in vitro* experiments. Collectively, the NOGEA was useful for identifying master genes that control disease initiation and co-occurrence, thus providing a valuable strategy for drug efficacy screening and repositioning. NOGEA codes are publicly available at https://github.com/guozihuaa/NOGEA.

## Introduction

The onset and progression of most complex diseases often involves the dysfunction of thousands of genes as well as certain altered interactions among them. High-throughput technologies such as gene expression profiling and whole genome sequencing have permitted the identification of an increasing number of disease associated genes (DAGs) [1], which may provide valuable insight into mechanisms of disease initiation and progression. However, as the existing DAGs are usually derived from multiple sources, they often contain large amounts of redundant or false positive information [2] due to collection bias and noise, such that causal relationships among these genes in most cases remain elusive. Therefore, identifying master genes that control disease state transitions from large numbers of DAGs plays a critical role in understanding disease initiation mechanisms. In addition, complex diseases show considerable comorbidity [3]. The master gene defects in one disease may initiate cascades of interactions that lead to the co-occurrence of multiple diseases in a given patient. Pharmacological targeting of the DAG module on the human interactome has proven to be a valuable strategy for drug efficacy screening [4]. At present, it is unclear whether the identification of master genes will further facilitate the network-based drug repositioning.

Recent trends in omics technologies and complex biological networks have led to a proliferation of attempts to find the master genes for different diseases. For example, genome-wide association studies (GWAS) have emerged as a powerful tool for detecting sequence variation associated with many human traits and diseases [5]. Due to the low-frequency of many mutations, GWAS usually require large cohort sizes to attain sufficient statistical power. More importantly, GWAS identify only the genetic risk factors associated with disease, rather than the master genes of the disease phenotypes because patient genomes contain a certain proportion of “passenger mutations” [6] and the initiation of many diseases is often triggered by the interplay between genetic and non-genetic factors. Transcriptome analysis is considered to be an effective complement of GWAS for its ability to capture non-genetic perturbations to the organism. Yet variations in mRNA expression are sometimes caused by aberrant protein activity of upstream regulators such as transcription factors, making it difficult to directly identify the master gene set using transcriptome profiling [7].

Recently, gene co-expression-based approaches have been proposed to construct context-specific regulatory networks [8] and a local network entropy measure has been developed based on co-expression networks for identifying master genes [9]. While these approaches provide new ways to find master genes, building a highly confident co-expression regulatory network often requires large sample sizes, which are usually not available for relatively rare diseases. To overcome this limitation, protein-protein interaction (PPI) network-based approaches have been developed to infer master genes that are important for disease-related biological processes, such as predicting therapeutic targets [10] or driver genes [11]. Some topological parameters such as the degree and betweenness centrality of the nodes are usually used as important measures to screen master genes [12]. However, current approaches are based mainly on the constant global undirected interactome, ignoring the fact that disease initiation and therapeutics are frequently context-dependent, depending on specific tissues or pathological microenvironment [13]. Therefore, some genes that exhibit important topological properties on the interaction network, such as the hub genes [14], will be automatically selected as key regulators for disease state initiation and maintenance ‘ leading to a possible increase in false positive master genes. Conversely, some classes of genes presenting as upstream regulators of a signaling cascade, such as the G protein-coupled receptors [15], may be identified as dispensable genes due to their relatively low degrees on the interactome, thus decreasing the sensitivity for distinguishing core ones from the giant pool of DAGs.

In this study, we have developed a network-oriented gene entropy approach to quantify the perturbation or regulatory ability of each DAG in distinct disease contexts by assembling and interrogating disease-specific regulatory networks. Master genes for each disease, whose altered expression was sufficient for disease state transitions, were identified as those genes that exhibited high entropy values by our *in silico* method, and were further adopted to investigate comorbidity and causal relationships among different diseases. We further confirmed that existing effective drugs are most likely to target the local module of master genes on the interactome. Using these methods, we have identified 11 old drugs as potent anticancer agents for pancreatic cancer treatment.

## Results and Discussion

### Computation of gene entropy in disease networks

To identify master genes in distinct disease contexts, a network-oriented gene entropy approach (NOGEA) was developed (Figure 1A and 1B). Briefly, Shannon entropy theory was applied to quantify the amount of disorder within intracellular signals in each disease specific context, which was subsequently factorized as the summation of contribution for each DAG. First, directed disease specific gene networks for 293 diseases were constructed to reflect the distinct disease contexts by mapping all DAGs (Table S1) to a previously established directed PPI network (Table S2) [20]. A directed network visualizes the hierarchy of intracellular signal transduction between the interacting proteins, and hence clearly reflects the importance of each DAG in a certain physiological and pathological context. The regulation likelihood between each pair of DAGs was then calculated based on the directed distance on the PPI network to generate a probability-based signaling flux matrix (Figure 1A). Finally, the perturbation ability of each DAG in a disease-specific context was calculated by the network-oriented gene entropy metric (Methods, Figure 1B). The distribution of entropy values for all DAGs is illustrated as a histogram in Figure S1, and the perturbation ability of each DAG was then ranked based on their entropy values (Table S1).

**Figure 1.**
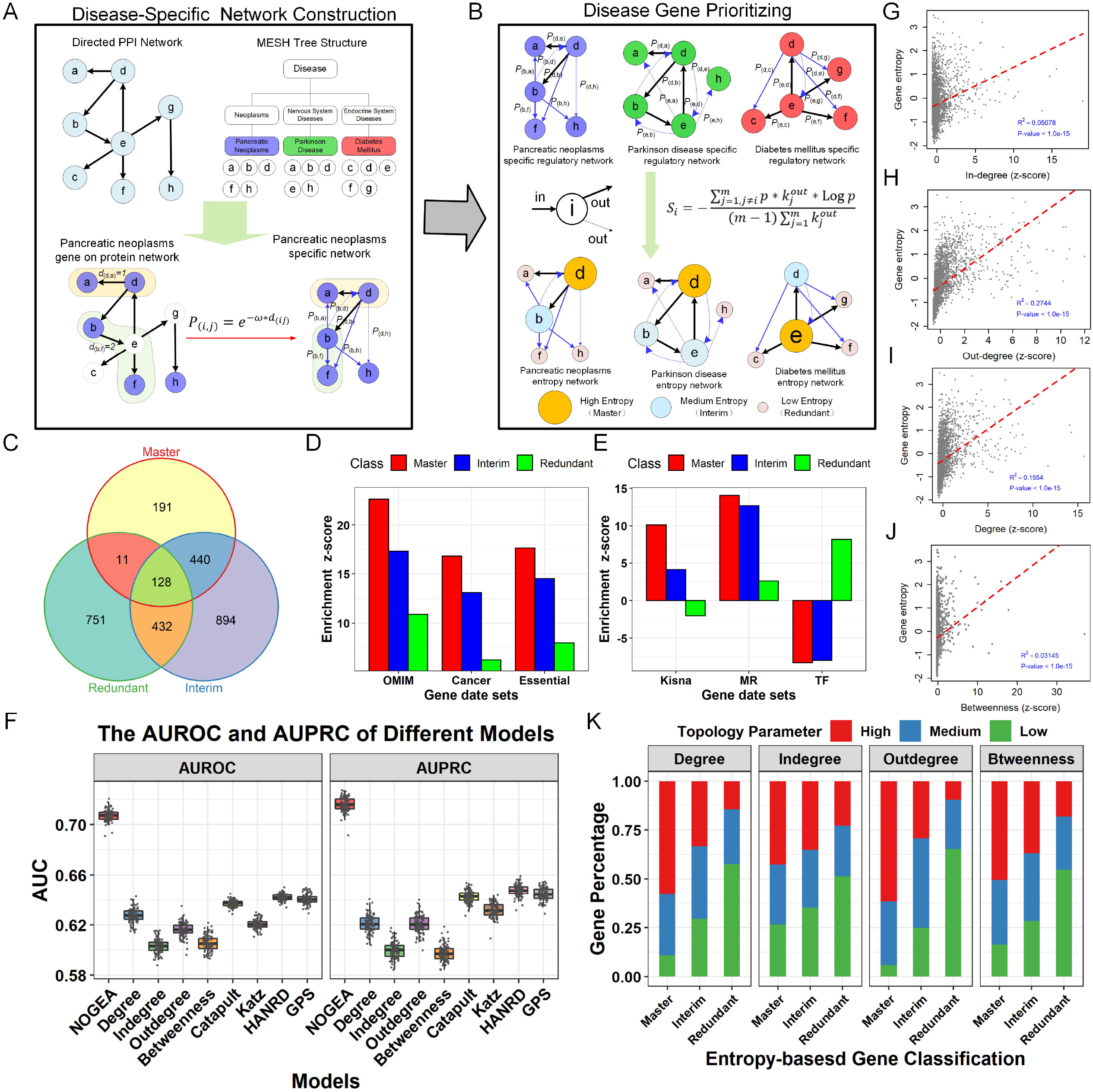
Computation and characterization of network-oriented gene entropy in disease-specific networks. A. Construction of directed disease-specific gene networks by mapping disease genes to the directed PPI network and normalizing the interaction strength. **B.** Calculation of the perturbation ability (gene entropy) of each gene. **C.** The Venn plot of the disease gene from different classes; Master: the master genes, Interim: the interim genes, Redundant: the redundant genes. **D.** Enrichment result (z-score) of master, interim and redundant genes in the context of OMIM, cancer and essential genes. **E.** Enrichment result (z-score) of master, interim and redundant entropy genes in the context of kinase, membrane receptor (MR), transcription factor (TF). **F.** Comparison of NOGEA performance with other methods for disease gene prioritization using AUROC and AUPRC. **G.** DAG entropy values versus their in-degree in the primary directed PPI network. **H.** DAG entropy values versus their out-degree in the primary directed PPI network. **I.** DAG entropy values versus their betweenness in the primary directed PPI network. **J.** DAG entropy values versus their degree (sum of in- and out-degree) in the primary directed PPI network. **K.** Assessment of the association between gene entropy and four commonly used network topology parameters.

To efficiently explore the biological features of each entropy distribution, all DAGs were classified as “Master”, “Interim” or “Redundant” genes which represent high, medium and low entropy genes, respectively. We created an entropy value curve for each disease and then identified two inflection points as thresholds to separate the low, medium and high entropy genes, respectively (Methods). We then merged the master genes of all diseases into a whole master gene set. Interim and redundant genes from different diseases were treated in the same way to obtain the whole interim and redundant gene sets, respectively. As a result, 798 master, 1,962 interim, and 1,387 redundant genes were obtained (Figure 1C, Table S3).

In order to verify whether the master genes play a key role in disease initiation and development, enrichment analyses were performed using several well-established gene clusters (Table S4). We observed that there was an overrepresentation (z-score=22.61) of disease-causing mutation-associated proteins among all master genes, which was higher than the enrichment score of both interim and redundant genes (Figure 1D). The essential genes were demonstrated to play critical roles in human diseases [28], and the master genes were enriched in essential genes, whose z-score was two times larger than the enrichment score of the redundant genes (Figure 1D). More importantly, we found that master genes were highly enriched in cancer-associated genes; whereas, redundant genes showed less enrichment (Figure 1D). Further KEGG analysis of the master genes showed that these genes were mainly enriched in pathways with close relationships with cancer initiation and progression (Figure S2). For example, PI3K-AKT signaling pathway (has:04151), which is commonly perturbed in cancers, were found among the top five enriched pathways (P < 10e-30). In a recent study, genes on the interactome were classified into different node types, in which “indispensable” nodes were found to be key players in mediating the transition of disease states. As shown in Figure S3A, we found that master genes were highly enriched in “indispensable” genes, but redundant genes were enriched among the “dispensable” genes. Consistent with these observations, the master genes were highly enriched in “critical” genes that acted as driver nodes in all control configurations (Figure S3B) [26]. Further dissection of all different functional classes within signaling proteins revealed that the master genes were most likely enriched in kinases and membrane receptors (Figure 1E). In summary, the results indicated that the master genes are preferred key regulators in disease initiation and development, reflecting the reliability of the NOGEA method.

Traditional network topology parameters, such as the connective degree and betweenness centrality, are commonly used as baseline methods for characterizing the importance of nodes in biological networks [29]. To validate the effectiveness of NOGEA, we compared it with four baseline methods (the connective degree, connective in-degree, connective out-degree and betweenness centrality-based methods) and four newly proposed methods (Katz [30], Catapult [30], HANRD [31] and GPS [32]), all of which are network-based methods for prioritizing disease genes. We first compared the AUROCs between different methods (Methods) and found that NOGEA significantly outperformed both the baseline methods and the newly proposed methods (Figure 1F). We further evaluated AUPRC, area under the precision-recall curve, for each method. NOGEA consistently surpassed all other methods, overmatching the second-best method by ~10% (Figure 1F).

Correlations between gene entropy values and four network topology parameters were assessed using Pearson’s correlation coefficients (PCC). For most diseases, we observed that the PCCs between gene entropy values and network topology parameters were relatively small (<0.25, Figure S4A). Nonetheless, significant correlation values were observed between the in-degree connective (R^2^=0.051, P<1.0e-15, Figure 1G), out-degree connective (R^2^=0.274, P<1.0e-15, Figure 1H) degree connective (sum of in and out-degree, R^2^=0.155, P<1.0e-15, Figure 1I) and betweenness centrality (R^2^=0.031, P<1.0e-15, Figure 1J) for genes in the primary directed PPI network versus gene entropy values. Fisher’s exact test was then applied to further determine whether gene entropy is associated with traditional network topology parameters. Specifically, we constructed a contingency table to classify the disease genes into different bins based on their entropy values and network parameter values (Figure 1K). We found that gene entropy was significantly associated with traditional network topology parameters, including connective degree (P < 0.01), connective in-degree (P < 0.01), connective out-degree (P<0.01) and betweenness centrality (P<0.01). All these results demonstrate that master genes prefer to possess high topology parameter values, indicating relative consistency between gene entropy and the four network topology parameters.

To investigate variation of the regulatory role of a specific gene in different diseases, we calculated the divergence-degree of gene entropy across diseases using the coefficient of variation (CV) (Table S1, Figure S4B). The results show that up to 60% of the genes have a high CV (>15%), indicating the distinct roles these genes play in different disease contexts. We then examined the entropy value variation of the shared genes in different diseases, and observed that these genes usually exhibit similar entropy values in distinct diseases within the same disease category. For example, corticotropin-releasing hormone receptor 1 (CRHR1) is related to eight mental health-associated diseases with different entropy rank scores (rank>0.80), including anxiety and depressive disorders (Table S1), which is consistent with its major role in mental disorders [33]. We also observed a low entropy rank score for CRHR1 in pulmonary disease (rank=0.55), indicating variation in its regulatory role in distinct disease contexts. Further, we found that ~15% genes have approximately equal rank scores in their associated diseases. For instance, interleukin 4 receptor (IL4R) and phosphatidylinositol-4,5-bisphosphate 3-kinase catalytic subunit alpha (PIK3CA) had high rank scores in their associated diseases (Table S1), especially for neoplasms, suggesting crucial roles for these genes in these diseases. In summary, NOGEA provided a new way to explore the regulatory role of each DAG in distinct disease contexts.

### NOGEA for exploring disease comorbidity

Exploration of the underlying mechanisms of comorbidity, which refers to the coexistence of multiple diseases or disorders, is difficult due to complex interactions among environmental, lifestyle and treatment-related factors [34]. In addition, disease comorbidity includes not only the co-occurrence of multiple diseases, but also the potential cause-and-effect relationships among these diseases. Thus, uncovering the diseases’ co-occurrence and causal relationships along with underlying mechanisms is of great significance for their prevention and treatment. Using experiment-based approaches or mathematical models, previous studies explored the molecular features of disease comorbidity for several diseases, including from gastritis to gastric cancer [35] and from diabetes to cancer [36]. However, existing experiment-based methods to explore the underlying mechanisms for co-occurrence and causal relationships remain costly, labour-intensive, or focused on a small fraction of molecular features. Comparatively, mathematical models provide novel ways to reveal disease comorbidity using multi-omics data; however, these models are difficult to apply in other diseases, due to the lack of multi-scale information for these diseases.

The results discussed above demonstrate that NOGEA-inferred master genes are closely associated with disease onset and development, prompting us to investigate whether the network entropy-based approach would be capable of uncovering the molecular basis of disease co-occurrence. Therefore, we constructed a new master gene disease network (M-GDN), where edge would link two different diseases if they shared at least one master gene (Table S5). For comparison, we constructed five other disease networks: the redundant gene-based disease network (R-GDN), the interim gene-based disease network (I-GDN), the all genes-based disease network (A-GDN), the traditional hereditary disease network (THDN) and the random disease genes network (RGN).

To test whether the M-GDN would provide an accurate picture of disease comorbidity, we evaluated the Tanimoto similarity between these networks and the human disease comorbidity network (HDCN), which was extracted from the Medicare Claims Database and constructed in a recent study [3]. The M-GDN showed the highest similarity with the HDCN (higher than that of R-GDN and THDN) at a significantly higher level than expected based on the random values (Figure 2A), which indicates that those genes most associated with disease comorbidity tended to be master genes with high entropy rather than arbitrary disease genes. In contrast to previous THDN models, M-GDN considers genetic factors as well as genes that respond to environmental, lifestyle, and/or treatment-related factors, thus providing a more comprehensive solution for exploring the comorbidity of disease. Furthermore, in view of the impact of cellular network interactions on disease comorbidity, we extended our result to a PPI-based M-GDN (Table S6), where two diseases were linked if the master gene of one disease directly interacted with genes of the other disease in the PPI network. Consistent with the above results, the PPI-based M-GDN demonstrated the best predictive ability in identifying disease comorbidity. We then observed that the inferred underlying molecular mechanisms of disease comorbidity are in accordance with current pathobiological knowledge (Figure 2B). For example, M-GDN confirmed the conclusion that AKT1 mutations lead to schizophrenia and type 2 diabetes mellitus (with rank scores of 0.96 and 0.94 in schizophrenia and type 2 DM, respectively) [37]. We also observed in the M-GDN that ADRB2 mutations may lead to asthma and obesity (with rank scores of 0.95 and 0.97 in asthma and obesity, respectively), which is consistent with a previous study [38]. These results suggest that M-GDN helps bridge the gap between bench-based biological discovery and bedside clinical solutions, and thus may provide new insights into the mechanisms of disease comorbidity.

**Figure 2.**
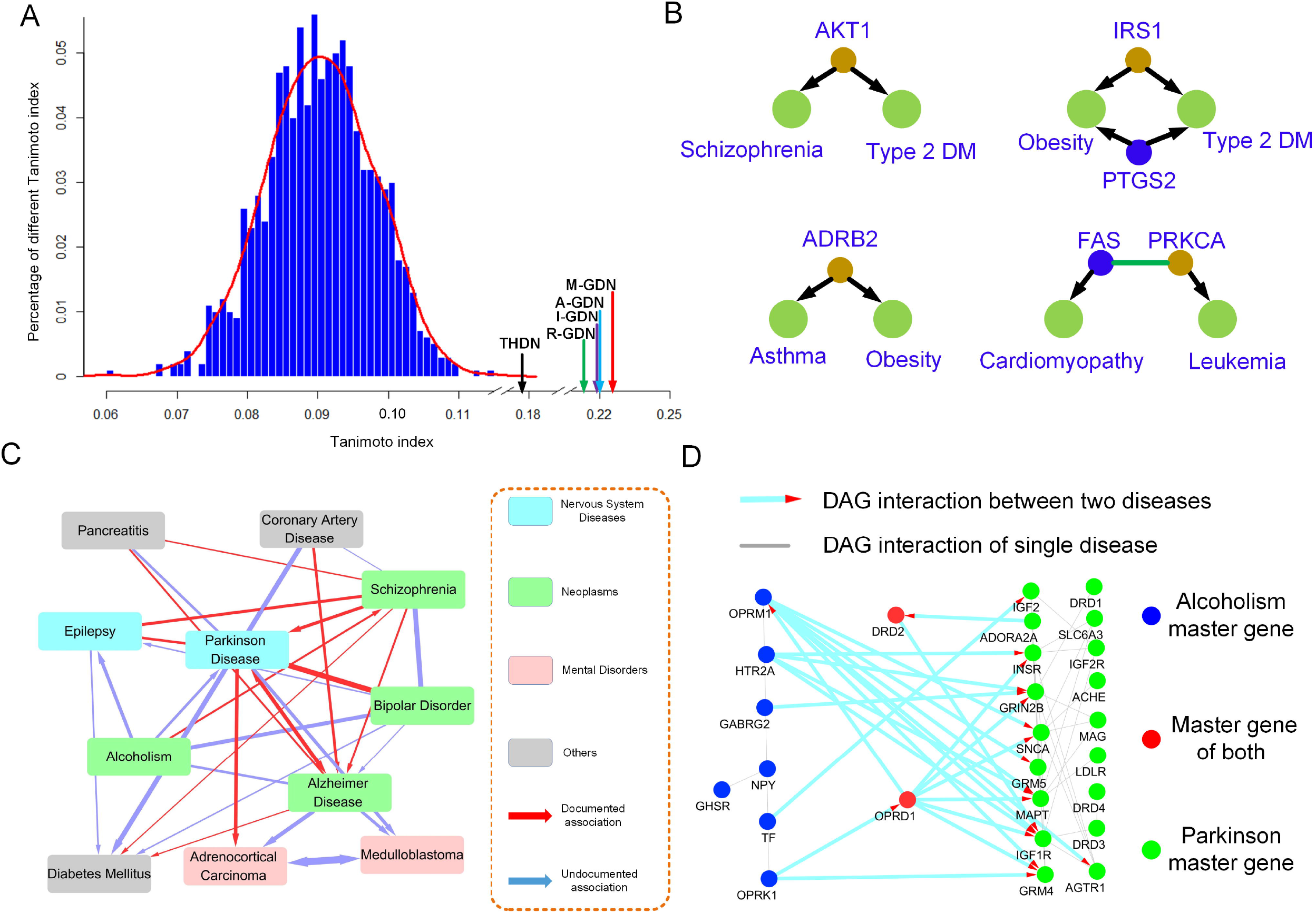
Exploration of disease comorbidity using network entropy. A. Distribution of Tanimoto similarities between HDCN and other disease-disease networks (M-GDN, I-DGN, R-DGN, A-DGN, THDN and RGN). **B.** The inferred molecular basis of disease comorbidity relationships. Brown and blue nodes represent master genes inferred by NOGEA; green nodes represent diseases. **C.** The comorbidity of Parkinson’s disease. In this figure, the width of the edge represents the likelihood of disease comorbidity, arrows represent the inferred causative disease-disease associations, and the color of the nodes depicts the disease category from MESH. **D.** The molecular basis of the comorbidity between Parkinson’s disease and alcoholism. The nodes represent the master genes of the disease and the directed links describe the direction from the directed PPI network.

Recent reports in the literature suggest that mutations in the IRS1 gene are closely related to the comorbidity of type 2 DM and obesity [39]. The M-GDN revealed that, in addition to IRS1, PTGS2 also plays a crucial role in the co-morbidities of these diseases. It is well known that PTGS2 influences the inflammatory response, which is closely connected with the comorbidity of type 2 DM and obesity [40]. Another example is the comorbidity of leukemia and cardiomyopathy, whose underlying mechanisms remain unclear. Interestingly, FAS is involved in the regulation of cell apoptosis, which affects left ventricular function [41] while PRKCA enhances cell resistance [42] and regulates cardiac contractility and an increased risk for heart failure. More importantly, the FAS-PRKCA interaction has been identified as the top connected cross-talk PPI by *in situ* proximity ligation assays [43]. These results demonstrate that the interaction between FAS and PRKCA may account for the comorbidity of leukemia and cardiomyopathy.

Next, we investigated the molecular basis of disease causal relationships from the perspective of directed biological networks. As an illustration, we constructed a directed comorbidity network (Table S7, Figure 2C) centered on Parkinson’s disease. We observed high co-occurrence risk between Parkinson’s and other diseases including Alzheimer’s disease. Recent research suggests that these diseases are related to the accumulation of common proteins in the brain, such as alpha-synuclein protein [44]. Using alcoholism and Parkinson’s disease as an example, we observed a significant directed interaction from alcoholism to Parkinson’s disease (P<0.01), but not vice versa. This result is consistent with recent clinical studies, which suggest that alcoholism may an inducer of Parkinson’s disease [45]. A subsequent network analysis further discovered that the aberration of alcoholism master genes may lead to the modification of most Parkinson’s disease’s master genes (Figure 2D). Collectively, NOGEA is potentially useful for investigating mechanisms underlying disease comorbidity as well as their causal relationships.

### NOGEA can infer drug-disease associations

Recently, several state-of-the-art network-based methods were proposed to investigate the relationships between drugs and diseases, such as the network proximity approach and network inference algorithm [4, 46]. In this study, we assessed relationships between DAGs and drug targets based on the gene network entropy to evaluate the effects of drugs on each disease. For each drug-disease relationship, we calculated the drug disturbance entropy (DDE) parameter, which represents potential therapeutic effects of the drug (Methods, Table S8-S10). To further investigate DDE’s effectiveness, we evaluated the correlation between the DDE value and the hits by known drug-disease interactions (DDIs), and found the occurrence number of known DDIs increased with increasing DDE values (Figure 3A). Consistent with previous research [4], a highly significant correlation occurred between DDE values and the enrichment of known drug-disease interactions (R^2^=0.75, P=2.2e-16) (Figure 3B), indicating a high likelihood that a drug will successfully treat a disease if the drug is capable of strongly perturbing the local module of master genes on the interactome.

To validate the utility of DDE for distinguishing known drug-disease pairs from the unknown drug-disease pairs, we compared the AUC of ROC curves for different drug-disease prediction methods (Methods). To obtain a robust AUC estimation, the drug-disease set was split into a training set and a testing set according to a given fraction coefficient for developing and validating the model, respectively. We compared the DDE’s performance with several other state-of-the-art methods [4, 46], including the network inference algorithm (NIA), network proximity approach (NPA), network kernel approach (NKA), network shortest approach (NSA), network center approach (NCA), and network separation approach (NSEA). As illustrated in Fig. 3C, DDE exhibited the best performance (average AUROC=70%) in discriminating known and unknown drug-disease pairs, significantly outperforming the other approaches. Interestingly, we noticed that the NIA, which appeared to be the second-best method (average AUROC=68%), was also able to construct a directed disease-specific gene network and identify master genes before predicting the drug-disease associations. A compressive comparison between the two methods demonstrated their connection and difference (as seen in Supplementary Note 2, Figure S5, Table S11-S12). Collectively, these results suggest that DDE is effective for predicting drug-disease associations.

**Figure 3.**
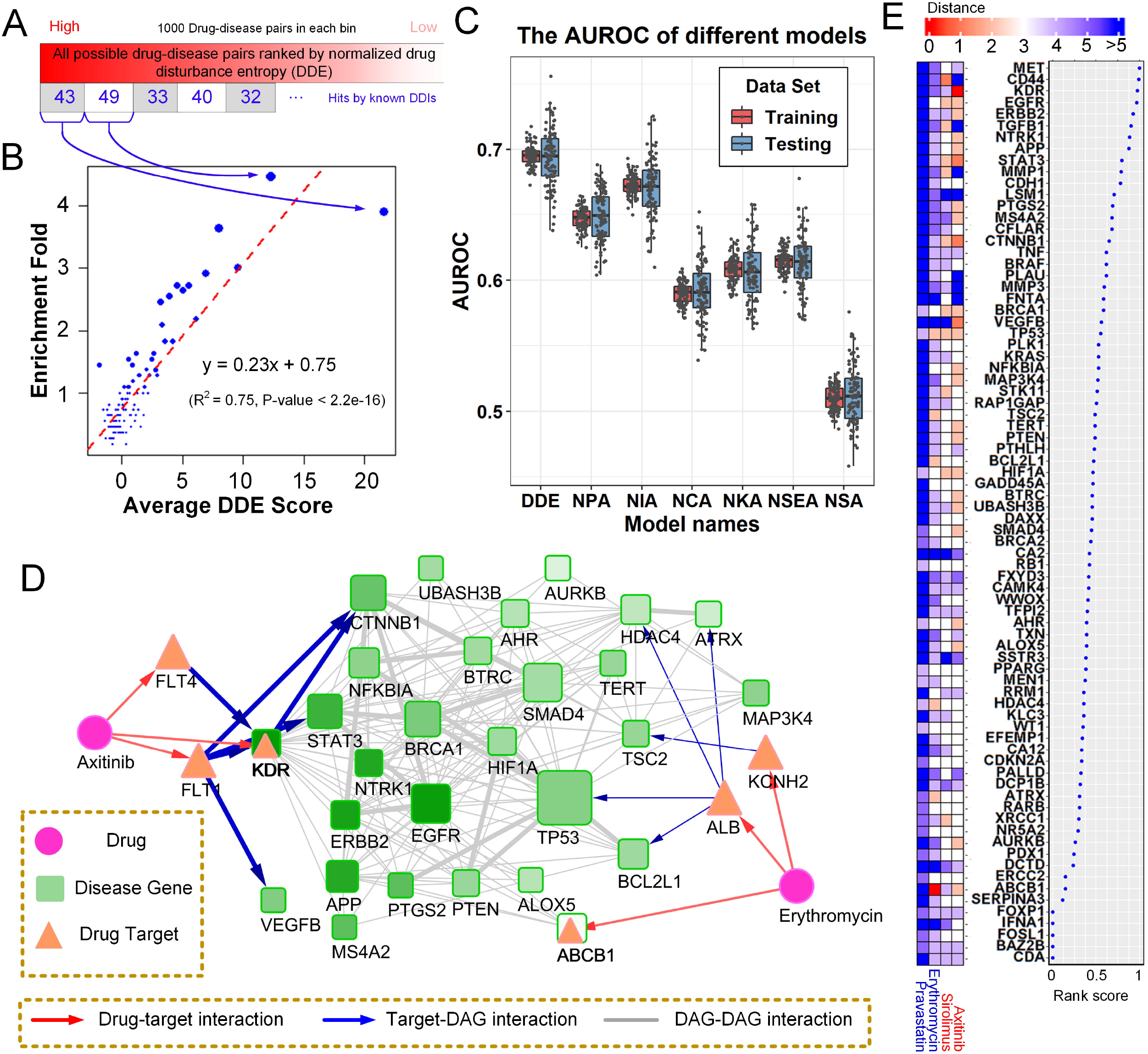
Drug-disease association inference based on the disease gene entropy. A. The hits number by known DDIs in each ranked drug-disease pair bin. **B.** The correlation between average DDE score in each bin and the hits enrichment fold for known DDIs. **C.** AUROC for drug-disease predictions using different methods. **D.** The interaction between drug targets and pancreatic cancer genes. The width of the links, the shade of the pancreatic cancer genes nodes, and the size of the node describe the interaction strength, entropy value, and degree of each node in the human interactome, respectively. **E.** The entropy value rank plot of pancreatic cancer genes (right); the heat map describes the shortest distance between the drug targets and pancreatic cancer genes of four drugs (left).

Pancreatic cancer is a refractory malignant carcinoma of the digestive tract with a 5-year survival rate of ~4% [47] that modestly responds to very few existing chemotherapy treatment options. Revisiting the complex interaction pattern between drug targets and pancreatic cancer genes in a systemic manner is essential for developing more effective therapeutic regimens. Therefore, we used pancreatic cancer as an example to explore the utility of NOGEA for drug-disease association inference. By measuring the entropy of each pancreatic cancer gene in the pancreatic cancer specific network (Figure 3D, Figure 3E), we found that those genes with high entropy such as MET, KDR, and EGFR may play more important roles than the lower entropy genes for pancreatic cancer treatment. As reported in previous studies [48], EGFR-mediated signaling is involved in the tumorigenesis of pancreatic cancer, and the preclinical data support EGFR inhibition as a potential treatment strategy for pancreatic cancer. In addition, c-Met protein, which is coded by the MET gene, is a marker of pancreatic cancer stem cells and thus a therapeutic target [49]. KDR (VEGFR-2) is known to be crucial for embryonic vasculature development by modulating endothelial cell proliferation and migration [50]. Moreover, the CD44 gene is a potentially interesting prognostic marker and therapeutic target in pancreatic cancer [51].

To investigate differences in the targeting patterns between effective drugs and other less-effective drugs from a network-based perspective, we constructed a gene entropy map for pancreatic cancer. We first calculated the linkage strength between drug targets and pancreatic cancer genes for two FDA-approved drugs: Axitinib and Erythromycin (Figure 3D). Axitinib binds to FLT4, FLT1 and KDR, which was identified as a pancreatic cancer master gene by NOGEA. The DDE of Axitinib to pancreatic cancer is 37.6, suggesting that targets of Axitinib are more closely related to pancreatic cancer genes than expected by chance. Conversely, the DDE of Erythromycin (whose efficacy remains unknown) to pancreatic cancer is 1.1. Even though this drug inhibits ABCB1, ALB and KCNH2, the disease proteins and drug targets are not closer than expected by randomly selecting protein sets. However, some drugs that do not directly inhibit the pancreatic cancer master genes may still have the potential to be effective drugs. For example, Sirolimus, which is currently in phase II clinical trials, targets three proteins (FKBP1A, FGF2 and MTOR) but no known pancreatic cancer genes. Nevertheless, Sirolimus has a high DDE value of 12.1 due to the relatively strong perturbation of high entropy genes such as CD44 and EGFR via FGF2 (Figure 3E). Drugs such as Pravastatin (DDE=-0.7) are predicted to be ineffective pancreatic cancer drugs due to their weak perturbation of nearly all pancreatic cancer genes (Figure 3E). Collectively, these results suggest that NOGEA may be capable of identifying the core genes among many DAGs that provide the basis for rational drug discovery.

### Pancreatic cancer drug screening

Due to the encouraging performance of the drug disturbance entropy metric for accurately inferring drug-disease associations, we screened potentially effective drugs for pancreatic cancer treatment. We first calculated and prioritized DDE values for all FDA-approved drugs (Table S13-S14). From top 10% of these drugs, we selected 19 molecules that were not known to be associated with pancreatic cancer for further experimental validation. The half-maximal inhibitory concentration (IC_50_) of a molecule, an important metric to measure its response to certain cancer cell lines, has been widely applied in the screening of potential anti-proliferative agents in preclinical cancer pharmacogenomics. The BxPC3 human pancreatic cancer cell line, which has been frequently used in the study of pancreatic cancer and screening of chemo preventive agents [52], was used in our *in vitro* study to evaluate its response to the candidate drugs. We identified 11 candidate drugs that inhibit BxPC3 cell lines in a dose dependent manner and exhibit low IC_50_ values (<100 μM/L, Figure S6, Figure 4A-4C), demonstrating their efficacies for inhibiting pancreatic cancer cell proliferation and potential for pancreatic cancer therapy *in vivo.* One drug for example, Vinorelbine, is a drug that has already been approved for non-small-cell lung cancer treatment [53]. In our study, Vinorelbine exhibited a low IC_50_ value of 1.55 nM/L (Figure 4A). Conversely, some non-classical anticancer drugs also displayed acceptable suppressive effects on BxPC3. Additional drugs, including Saquinavir, which is mainly used with other medications for HIV/AIDS treatment or prevention [54], and Celecoxib, a drug mainly used for treatment of pain and inflammation in adults [55], showed IC_50_ values of 22.63 μM/L (Figure 4B) and 45.36 μM/L (Figure 4C), respectively. These results indicate that our model has the capacity to predict proper drug candidates for disease therapy.

**Figure 4.**
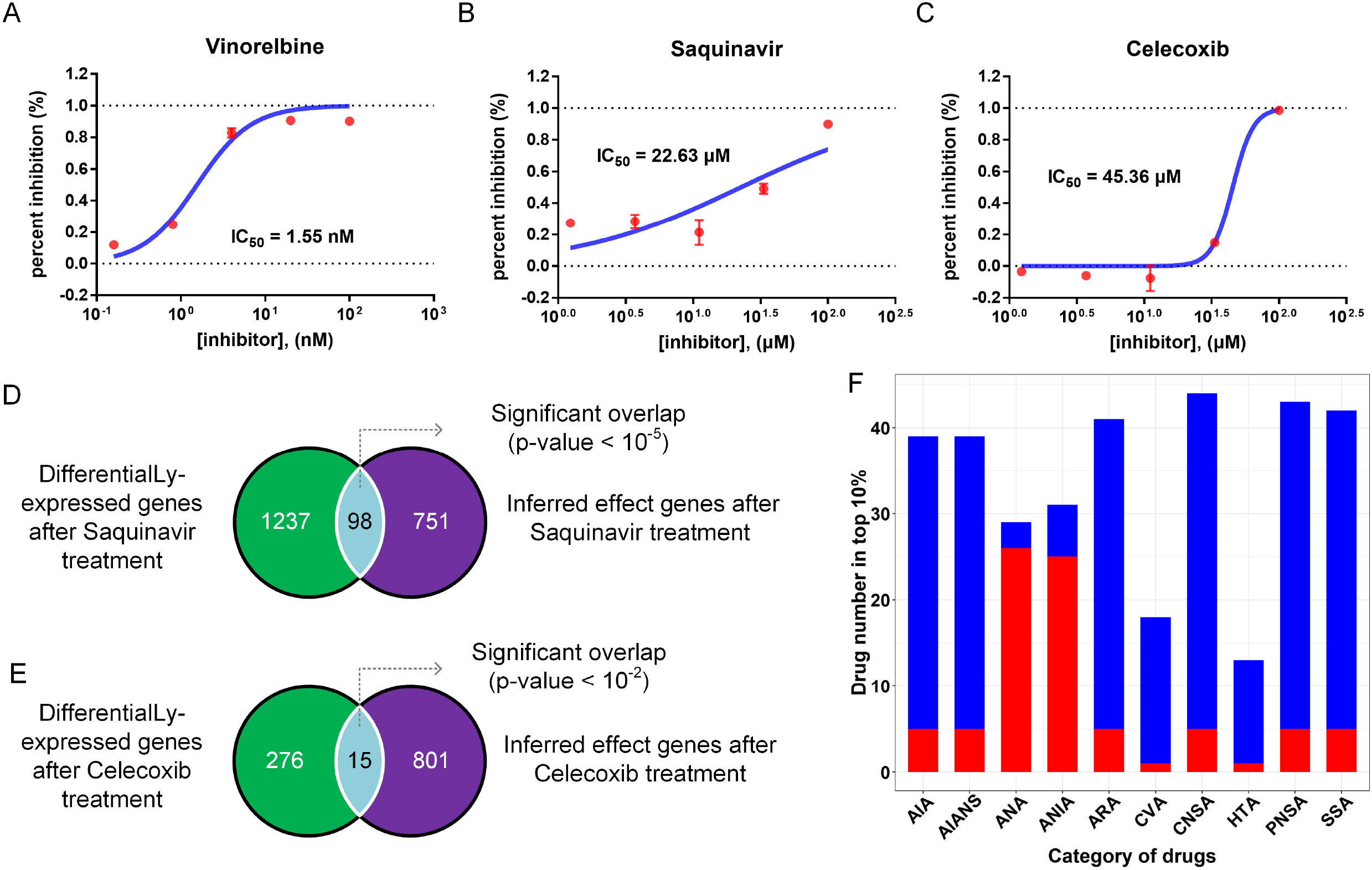
Screening of potential efficient drugs for pancreatic cancer treatment. **A-C.** Cell inhibition rate curves against BxPC3 for Vinorelbine, Saquinavir and Celecoxib, respectively. **D.** The number and significance of overlapped genes between differentially expressed genes and the inferred effect genes after Saquinavir treatment. **E.** The number and significance of overlapped genes between the differentially expressed genes and the inferred effect genes after Celecoxib treatment. **F.** The overlapped drug number between each category and the top 10% of efficient drugs. Red bar: number of literature mining significant drugs; AIA: Anti-Inflammatory Agents, AIANS: Anti-Inflammatory Agents (Non-Steroidal); ANA: Antineoplastic Agents; ANIA: Antineoplastic and Immunomodulating Agents; ARA: Antirheumatic Agents; CVA: Cardiovascular Agents; CNSA: Central Nervous System Agents; HTA: Hypotensive Agents; PNSA: Peripheral Nervous System Agents; SSA: Sensory System Agents.

Transcriptional expression analysis was conducted to validate our hypothesis that efficient drugs tend to perturb the master genes directly or through their targets. We first identified 1,335 differentially expressed genes (referred to as SAQDEGs) after Saquinavir treatment (Figure S7A, Table S15). The pancreatic cancer master genes (n=849) that were most likely to be perturbed by Saquinavir were named SAQPEGs and further incorporated with their corresponding neighbor genes on the interactome (Table S15). Finally, a hypergeometric test was used to assess the overlap between SAQDEGs and SAQPEGs. These analyses revealed that the differentially expressed genes were significantly enriched for SAQPEGs (Figure 4D, P<0.01). Results for Celecoxib were similar to those for Saquinavir (Figure S7B, Figure 4E), suggesting a close relationship between genes perturbed by the efficient drugs and the local module of master genes.

Finally, to demonstrate the reliability of the DDE approach for extensive screening of pancreatic cancer candidate drugs, we conducted a literature mining analysis to evaluate the association between the candidate drugs (top 10%) and pancreatic cancer based on our previous reports [56] (Methods). We observed that 8 of the top 10 candidate drugs were anticancer agents that showed significant literature mining correlation scores with pancreatic cancer (P<0.01, Table S14). In addition, most anticancer candidate drugs (~85%) were significantly associated with pancreatic cancer (Figure 4F, Table S14), suggesting the sensitivity of this model. Interestingly, an analysis of the categories of these candidate drugs revealed that the largest proportion, 44/224 (19.6%), were assigned to Central Nervous System Agents (CNSA). For example, Celecoxib, which was sensitive to the BxPC3 cell lines as mentioned above (Figure 4C), also acts as a CNSA. In general, these results indicate that DDE provides a rational strategy for drug repurposing due to its capacity to quantify drug targeting tendencies on the interactome.

## Materials and methods

### Data set collection

The DAGs for all diseases were obtained from four publicly available databases including KEGG Disease [16], Comparative Toxicogenomics Database [17], Therapeutic Target Database [18] and PharmGKB [19]. All disease names and their corresponding IDs were standardized by mapping to Medical Subject Headings ontology (MeSH; www.nlm.nih.gov/mesh/) and official gene symbols for these DAGs were retrieved from GeneCards (http://www.genecards.org/). We then conducted a disease filtering process to ensure disease specificity. We first removed diseases with levels < 2 on the MeSH tree structures, such as “Nervous System Diseases” and “Cardiovascular Diseases”, as these disease types are too broad. Tanimoto similarity (ratio between the number of shared DAGs and the number of joined DAGs) was then computed for each disease pair and used to remove diseases showing high similarity (>0.50) with its descendant disease. The weighted directed PPI network was constructed using data from a previous study [20], which consisted of 13,684 weighted interactions among 6082 proteins. The DAGs were then mapped to corresponding proteins in the PPI network, and those diseases with at least 20 DAGs in the human interactome were retained, for they are likely to induce a module on the network. As a result, we obtained 11,414 disease-gene associations between 274 diseases and 2848 protein-coding genes. For each disease, we manually extracted drug-disease associations from the drug indication information in DrugBank [21]. In addition, we obtained drug-target interactions for all FDA-approved drugs from DrugBank. To construct a disease comorbidity network, we retrieved disease pairs with comorbidity relationships from a recent study [3] of 665 diseases and their corresponding genes extracted from Online Mendelian Inheritance in Man (OMIM) [22].

### The disease-specific network-oriented gene entropy approach (NOGEA)

#### Construction of a flux matrix based on the expectation of the Bernoulli distribution

To construct the directed disease-specific gene networks, DAGs were mapped to the directed PPI network. For any given disease *D*, whose *m* associated genes can be mapped to the directed PPI network, an initial DAG vector 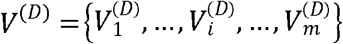 was generated to represent the disease, where 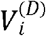 is the 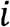-th DAG. The directed shortest path between two DAGs of disease *D* was calculated using the “igraph” package [23] based on the R 3.32 environment (r-porject.org). For a given DAG pair 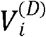 and 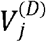, *I*_(*i, j*)_ is a random variable that obeys the Bernoulli distribution and represents the interaction or information transfer between node pair 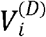 to 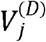. The distribution function of *I*_(*i, j*)_ is defined as

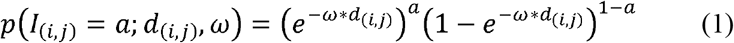

where *a* = 1 or 0, indicating whether signal transduction exists between node pair 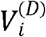 and 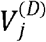, and *ω* is a scale parameter to adjust the likelihood for different distances. In addition, *d*_(*i,j*)_ is the directed distance between the given node pair 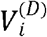 and 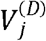. It is the number of edges in a directed shortest path connecting them, and was calculated using the “igraph” package based on Dijkstra’s algorithm, reflecting the possibility of the pairwise regulatory relationship from 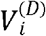 to 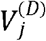. The details for determining the optimal scale parameter are presented in Supplementary Note 1. Therefore, the space of “possible” values assumed by *I*(*i, j*) is {0,1}, and if *a* = 1, *p*(*a; d*_(*i, j*)_, *ω*) represents the likelihood that there is a signaling flux between the node pair. In the field of network communication, it is widely accepted that the success rate of signal propagation decays exponentially with increasing distance [24]. In addition, previous studies have demonstrated that exponential decay is a popular kernel to characterize the network influence between two nodes [25]. Previously, we used the exponential component to evaluate the association between two nodes in protein-protein networks [26]. Thus, we believe that the success probability of the signal transduction between two proteins decays exponentially with the increase of their distance and the exponential component *e*^−*ω*d*_(*i,j*)_^ is useful for representing the success probability. In this way, the stochastic information flux matrix for a given disease is obtained by a simplified formula Eq. (2)

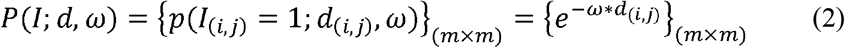

And, *p*(*I*_(*i, j*)_ = 1; *d*_(*i, j*)_, *ω*) is equal to the expectation of *I*_(*i, j*)_, where

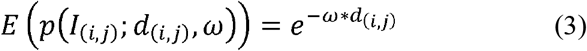

The expectation was subsequently used to estimate the distribution of signaling fluxing. For a given disease D with *m* associated genes, the biological signaling may flux between any node pair (DAG) 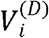 and 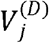. We then assumed that the edge (or the node pair) through which the signals fluxes is a random variable *F*, and its event space is

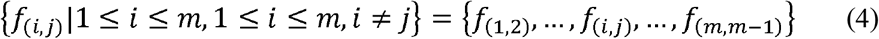

where *f*_(*i, j*)_ represents signals that may be transferred from DAG 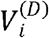 to 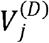.

#### Normalization of the fluxing matrix

The probability distribution of signal fluxing was estimated from

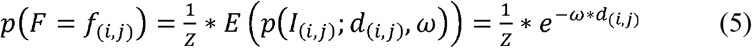

where *Z* is the normalization constant or partition function, and

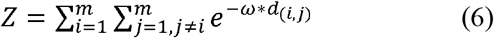

to ensure that the sum of the probability is 1.

#### Definition and calculation of disease gene entropy

Based on the probability distribution of signal fluxing, we calculated the entropy for a given disease *S*^(*D*)^ in terms of the weighted Shannon entropy formula, which can be interpreted as the degree of disorder or complexity for the disease specific context,

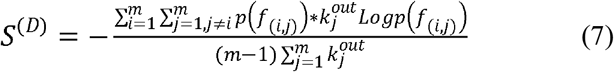

where 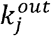 is the out-degree of node 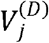 in the directed PPI network, which was calculated using the “igraph” package. Interestingly, we found that the disease entropy *S*^(*D*)^ can be factorized as shown in Eq. (8),

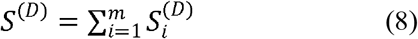

where 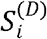 is the gene entropy of gene 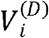, which is obtained by

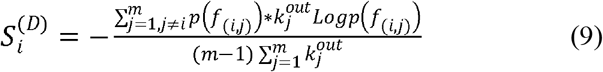

Therefore, 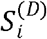 is a sub-entropy of disease entropy *S*^(*D*)^, and is considered as the “disorder contribution” to a disease specific context.

#### Gene entropy value normalization

Through the above procedure, a gene entropy map was established for 293 diseases. For any given disease *D*, the gene entropy z-scores were calculated, making the gene entropy values of different diseases comparable,

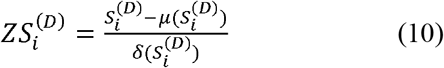

where 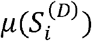 and 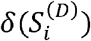 are the estimation of the expectation and standard deviations of 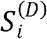 for disease *D*. In addition, to assess the disturbance capability of a gene in a disease-specific network in a more intuitive manner, we calculated the rank score for all DAGs according to their entropy values, which range from 0 to 1 and reflect their likelihood as master genes.

#### Rank score calculation of gene entropy

The gene entropy values for disease *D* were sorted in an ascending order, and a rank list was generated:

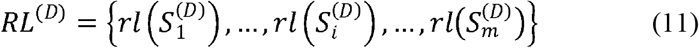

where the 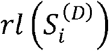 is the rank value of 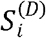 Note that those genes that possess equal entropy values have the same rank values. For example, if there are *k* genes 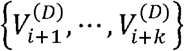 possessing equal entropy values their rank values 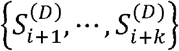, were determined by equation (12):

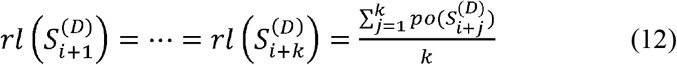

where 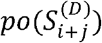 is the position of 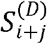 in the ascending entropy value list. Based on the rank list, rank score vector *RS*^(*D*)^ was generated by Eq. (13):

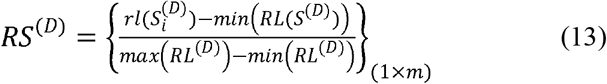

where *max*(*RL*^(*D*)^) and *min*(*RL*^(*D*)^) are the maximum and minimum of *RL*^(*D*)^, respectively.

#### Disease-gene classification based on the gene entropy value

To comprehensively explore the biological meaning of the entropy, we divided all DAGs into three groups based on their entropy values using an adaptive approach. Briefly, we created an entropy value curve for each disease, and identified two inflection points in the curve as thresholds. Specifically, for each disease D, we ranked each gene entropy value 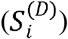 in ascending order. Then we mapped each entropy value onto a two-dimensional coordinate system such that the lowest entropy value 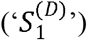 became coordinate 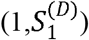, the second lowest value became 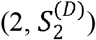, and so on, until the maximum entropy value 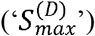 was reached. Two inflection points, individually defined as the threshold points of most rapid increase from the low to the medium and from the medium to the high entropy values, were identified in the entropy value curve from the interval of 10th to 50th percentile and 51st to 90th percentile, respectively, of all entropy values. The entropy value corresponding to this threshold was used as an adaptive disease-specific classification threshold. Master genes of all diseases were then merged and adopted as the whole master gene set to explore their common biological meanings. Interim and redundant genes from different diseases were treated in the same way to obtain the whole interim and redundant gene sets, respectively. Therefore, some genes may belong to all three gene sets (master, interim and redundant), because they play different roles in distinct disease contexts.

### Disease comorbidity relationship evaluation

A real human disease comorbidity network (HDCN) was constructed in which nodes represented diseases and edges represented the reported comorbidity relationships, respectively. We then built five different types of inferred disease comorbidity networks to compare with the HDCN. First, a master gene disease network (M-GDN) was constructed, where edges linked two different diseases only if they shared at least one high entropy gene. We then constructed the redundant gene disease network (R-GDN), the interim gene disease network (I-GDN), the whole genes-based disease network (A-GDN) and the traditional hereditary disease network (THDN), respectively. A Tanimoto coefficient was used to evaluate the similarity between different networks as shown in Eq. (15),

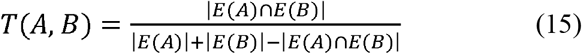

where *A* and *B* are different networks, *E*(·) represents the edge set of a given network and |*E*(·)| is the number of edges in the net. To assess the significance of the similarity of different networks, the random disease genes network was randomly generated 1,000 times and compared with the HDCN using equation (15). In the random disease genes network, each disease involves a random sampling gene set of the same size as the disease in A-GDN.

Previous research has demonstrated that cellular interaction links result in statistically significant comorbidity patterns [3]. Therefore, we believe that the directed interaction strength from the DAGs of one disease to another in the directed cellular network can reflect the causal relationship between the two diseases. To evaluate whether a causal relationship exists between two diseases, we estimated the significance of the interaction strength between the DAGs of the disease pairs using the Monte Carlo method. We first defined a raw causal relationship score (RCRS) for two given diseases: D1 and D2,

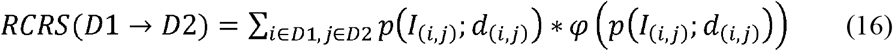

where *p*(*I*_(*i, j*)_;*d*_(*i, j*)_) was calculated by equation (1), *d*_(*i, j*)_ is the directed distance between master gene pair 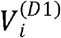 and 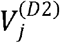, and *φ*(*p*(*I*_(*i, j*)_;*d*_(*i, j*)_)) is an indicator function. In addition, *φ*(*p*) was calculated as

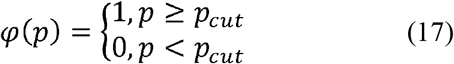

where *p_cut_* is a threshold, below which the probability was discarded and considered not contributive to the overall interaction and *p_cut_* was determined according to a previous study [27]. We then used a normalized causal relationship score (NCRS) to quantify the risk that disease *D*1 will induce disease *D*2. The *NCRS* is defined in Eq. (18)

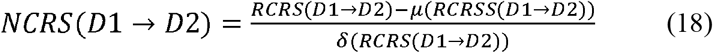

where *μ*(*RCRS*(*D*1 → *D*2)) and *δ*(*RCRS*(*D*1 → *D*2)) are the estimation of the expectation and standard deviations of *RCRS* under the same condition, respectively. Then, Monte Carlo simulation was performed 1,000 times to estimate the *μ*(*RCRS*(*D*1 → *D*2)) and *δ*(*RCRS*(*D*1 → *D*2)) by randomly sampling the same number of genes as *D*1 and *D*2. In each simulation, the values, the average and standard deviations of *RCRS* were calculated. To assess whether the causal relationship from disease *D*1 to *D*2 was significant, the P-value of *RCRS*(*D*1 → *D*2) was further calculated as shown in Eq. (19):

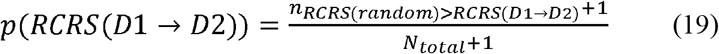

where *N_total_* is the total number of simulations, and *n*_*RCRS*(*random*)>*RCRS*(*D*1 → *D*2)_ is the number of random *RCRS* values that are larger than *RCRS*(*D*1 → *D*2). The *RCRS* value for the significance of P-values was set to 0.01. Finally, for a disease pair D1 and D2, if both *RCRS*(*D*1 → *D*2) and *RCRS*(*D*2 → *D*1) were significant (P<0.01), the two diseases were considered to be co-occurrent; whereas, if only one was significant (P<0.01), we determined that a causal relationship exists between the two diseases.

### Drug disturbance entropy (DDE)

To quantify the effects of a drug on each disease based on the gene network entropy, we applied an ensemble approach, referred to as drug disturbance entropy (DDE), to evaluate the relationship between drug targets and disease proteins (encoded by disease genes) on the interactome. We first evaluated the linkage strength between each DAG and drug target on the interactome, which was then transformed to a probability. The perturbation value for each target and DAG was defined as the product of the strength probability and the DAG entropy,

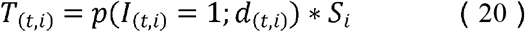

where *p*(*I*_(*t, i*)_ = 1; *d*_(*t, i*)_) represents the strength probability between drug target *t* and DAG 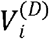, *S_i_* is the entropy value of DAG 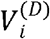, and *d*_(*t, i*)_ is the distance between target *t* and DAG 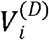. The raw disturbance entropy, which represents an estimate of a drug’s therapeutic effects through distinct targets, was defined as

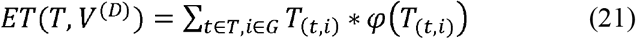

where *T*_(*t,i*)_ is the perturbation entropy between target *t* and DAG 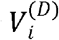, and *φ*(*T*_(*t, i*)_) is an indicator function as shown in Eq. (22)

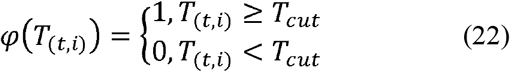

where *T_cut_* is a cut-off threshold of the disturbance entropy. The threshold of the perturbation value was determined by extensive sampling, and relationships with a perturbation value below this threshold were discarded. The remaining values were summed as the raw DDE of the drug to the disease. The advantage of this procedure is that weak relationships are eliminated, which greatly reduces noise and improves the robustness of the measure. By sampling across the range of *T_cut_* choices, the threshold that led to the highest ROC AUC was chosen. We obtained the proper *T_cut_* as 0.89 * *max*(*T*_(*t, i*)_) by evaluating the performance of predictions of drug-disease associations. Detailed information for determining *T_cut_* is depicted in Supplementary Note 1.

To avoid possible high DDE that may be caused by a large number of drug targets and DAGs, we converted raw DDE to a size-bias-free value using the mean and standard deviation of raw DDE modeled from sets of random molecules, so that the potential therapeutic effects between distinct drugs and diseases could be evaluated under the same metric. The raw drug disturbance entropy was transformed to a size-bias-free score under formula (23)

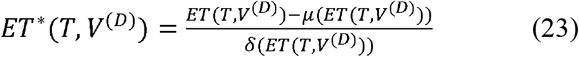

where *T* and *V*^(*D*)^ are the drug target set and the disease-associated gene set respectively; *μ*(*ET*(*T, V*^(*D*)^)) and *δ*(*ET*(*T, V*^(*D)*^) are the estimation of the expectation and standard deviations of DDE under this condition, respectively.

The estimation procedure of *μ*(*ET*(*T, V*^(*D*)^)) and *δ*(*ET*(*T, V*^(*D*)^)) are as follows: For each pair of (*T, V*^(*D*)^), we constructed 1,000 random set pairs with |*T*| targets and |*V*^(*D*)^| DAGs, preserving the degree distribution of the randomized targets and disease proteins. To avoid repeatedly choosing the same nodes during the degree-preserving random selection, we used a binning approach as described in a previous report [4].

## Conclusion

Disease phenotypes typically result from interactions among multiple complex environmental and genetic factors. The occurrence, development and treatment of a disease usually involves hundreds of genes [29]. Presently, we proposed a network-oriented gene entropy approach (NOGEA) for accurately inferring master genes that contribute to specific diseases by quantitatively calculating their perturbation abilities on directed disease-specific gene networks. Our results confirm that that master genes are enriched in gene sets that account for disease onset and development. This may imply that at a molecular level, those master genes with high entropy values are the underlying start-points of the disease state, impacting those redundant genes with low entropy through a directed disease-specific gene network. Interestingly, the comorbidity prediction model built using the master genes showed the best agreement with the independent clinical data set compared to the model established using the whole disease gene set. This indicates that our method may decrease the influence of noise and improve the efficiency for extracting more important genes from massive genomic data sets. Finally, through this method, 11 old drugs were newly identified and predicted to be effective for treating pancreatic cancer and then validated by *in vitro* experiments. However, it remains challenging to simulate the complex contents of the tumor microenvironment *in vitro*, making it difficult to comprehensively evaluate drug response using IC_50_. Therefore, despite our encouraging results, future work focusing on *in vivo* validation before clinical use is needed.

Although the identified master genes may be important for elucidating mechanisms of disease progression and drug screening, we acknowledge that it is difficult to directly evaluate the accuracy of NOGEA for identifying master genes at this stage due to the lack of ‘gold standard’ reference data sets. Nevertheless, the availability of more personal genome data in the future will allow for construction of patient-specific networks, NOGEA will provide new opportunities to identify patient-specific master genes and promote the development of personalized medicine. Emerging deep learning methods may become powerful techniques for exploring poly-pharmacy side effects [57] and discovering disease gene associations [58] from massive data sets [59]. Because gene entropy values can be used as novel disease feature data, we expect that integrating deep learning with NOGEA will significantly improve the accuracy for determining disease-drug or disease-disease associations. Extending the systematic approach presented here from signal drugs to multiple drugs may pave the way toward a better understanding of drug combinations.

## Authors, contributions

YHW and WX formulated the idea of the paper and supervised the research. ZHG and CH performed the research and drew the figures. XTC and SG collected data. ZYW and XTC performed laboratory experiments. ZHG, YXF, CH and CLZ wrote the paper. YXF, MS, PFT, YHM, JBZ, YL and ZHW revised the paper. All authors reviewed the manuscript.

## Competing interests

The authors have declared no competing interests.

## Acknowledgements

The research was supported by the National Natural Science Foundation of China (NO. U1603285) and the National Science and Technology Major Project of China (grant number 2019ZX09201004-001).

We thank TopEdit (www.topeditsci.com) for its linguistic assistance during the preparation of this manuscript.

## Supplementary material

**Figure S1 Distribution of gene entropy values for all DAGs**

Histogram plots showing the distribution of gene entropy values for all DAGs before (left) and after (right) normalization. The x-axis shows the range of gene entropy values, and the y-axis shows the count of genes possessing different entropy values.

**Figure S2 KEGG pathway enrichment results**

X-axis: the top 20 significantly enriched ‘KEGG pathway terms’ of the master genes; y-axis: significance of the enrichment [-log(P-value)].

**Figure S3 The disease-gene enrichment analysis for different classifications**

Enrichment results (z-score) of master, interim and redundant genes in the context of gene sets for critical (**A**), redundant (**A**), indispensable (**B**) and dispensable (**B**) genes.

**Figure S4 The property of the disease-gene entropy concept**

A. The correlation between entropy value and topology property for each disease. In this figure, each point represents a disease. The coordinate of each point represents the Pearson’s correlation coefficient (PCC) for the gene entropy values versus the in-degree (x-axis) and the out-degree (y-axis) of the disease-associated genes (DAGs). The size and the color represent PCC for the gene entropy values versus degree (sum of in- and out-degree) and betweenness, respectively. **B.** The distribution and cumulative probability of the coefficient of variation for the DAGs among different disease contexts.

**Figure S5 Rank scores for the top 20% of high entropy genes for three diseases**

Bar plots show the rank scores of the top 20% of high entropy genes for systemic lupus (CD4 cells) (top), systemic lupus (B cells) (middle) and rheumatoid arthritis (B cells) (bottom). Red bars represent the rank scores of the core genes retrieved from NIA.

**Figure S6 The dose–response curve of the BxPC3 cell of 8 drugs**

**A-H.** The dose–response curve of BxPC3 cells for 8 drugs that have not been associated with pancreatic cancer. X-axis: the concentration of each drug; y-axis: the percent inhibition rate of the BxPC3 cells.

**Figure S7 The heat map of microarray experiment results**

A. Differentially expressed genes between the Saquinavir (saq1, saq2) treated BxPC3 cell group and the control group (con1, con2). Color represent the relative expression of the differentially expressed genes. **B.** Differentially expressed genes between the Celecoxib (cel1, cel2) treated BxPC3 cell group and the control group (con1, con2).

**Figure S8 Estimation of the scale parameter ω**

Selected parameters (ω=1.1) that showed the highest mean AUROC and were thus used for further analysis.

**Figure S9 Characterization of gene entropy features with different scale parameters ω**

A. Normalized probability of different distances with scale parameter ω ranging from 0 to 4. **B.** Coronary disease gene entropy values with different scale parameters, ω=0 (top) and ω=10 (bottom). **C.** Coronary disease gene entropy values with scale parameter ω ranging from 0 to 10.

**Figure S10 Performance of the drug-disease relationship predictions using different scale parameters**

The box plot shows the AUROC for drug-disease predictions using different scale parameters. To account for the heterogeneous degree distribution of the directed interactome, we preserved the degree of randomized targets and disease genes.

**Table S1 Full list of disease-gene associations used in this study**

Entropy value: the entropy value calculated using NOGEA in a specific disease; rank score: the rank score for each gene entropy in a specific disease. This table also includes topology parameters of the DAGs in the directed global PPI network, i.e., the undirected degree, the in-degree, the out-degree and the betweenness centrality. In addition, this list includes the mean and standard deviations of the entropy among different diseases for a disease gene, the number of the gene-associated diseases and the coefficient of variation of the disease gene among different diseases. The evidence for the disease-gene associations was retrieved from CTD, TTD and PharmGKB.

**Table S2 List of the directed protein-protein interactions**

The list was obtained from a recent study as described in the paper, and each row presents a directed edge.

**Table S3 Classification of the disease-associated genes**

This list includes all the disease-gene relations used in this study. Genes of each disease were assigned to master, interim and redundant groups according to their entropy values.

**Table S4 Gene sets used for enrichment**

This table lists all 8 different gene sets used for enrichment analysis, which contains 1707 OMIM genes, 2186 predicted cancer genes, 1750 essential genes, 1551 transcription factors, 366 kinases, 249 membrane receptors, 1336 druggable genes and 982 FDA targets, respectively. All gene sets were obtained from a recent study (PMCID: PMC4983807).

**Table S5 Inferred comorbidity relationships of disease pairs from the shared genes**

This table lists all inferred comorbidity relationships involving master genes. As described in the paper, if two diseases shared a master gene, they were considered to be co-morbid diseases. Shared master genes are also listed.

**Table S6 Inferred comorbidity relationships of disease pairs from the interacting gene pairs**

This table lists all inferred comorbidity relationships involving master genes. As described in the paper, if master genes of two diseases directly interact with each other on the interactome, they were treated as co-morbid diseases. Interacting master gene pairs are also listed.

**Table S7 Inferred causal or co-occurrence relationships between Parkinson’s and other diseases.**

Results of the inferred relationships correspond with Figure 2C. This table lists all inferred causal or cooccurrence relationships between Parkinson’s disease and other diseases. The validated relationships are marked as “YES”. The “positive sim” is the likelihood from “V1” to “V2” and the “negative sim” is the likelihood from “V2” to “V1”.

**Table S8 Information for all FDA-approved drugs that were used in the present study**

This table lists all FDA approved drugs that were used in the present work and their corresponding IDs in other databases.

**Table S9 List of drug-target relationships used in the present study**

This table lists all FDA drug-target relationships used in this study.

**Table S10 Drug-disease information**

This table includes FDA drug indications, drug names and corresponding MESH IDs inferred from the indication information.

**Table S11 Gene rank list for three diseases**

This table lists the gene rank scores and core genes for systemic lupus (CD4 cells), systemic lupus (B cells) and rheumatoid arthritis (B cells).

**Table S12 Drug disturbance entropy (DDE) for each FDA-approved drug associated with three diseases**

This table lists the value of DDE calculated using NOGEA for each FDA-approved drug associated with the systemic lupus (CD4 cells), systemic lupus (B cells) and rheumatoid arthritis (B cells).

**Table S13 FDA-approved drugs and their categories**

This table lists all present FDA approved drugs and their corresponding categories retrieved from the DrugBank database.

**Table S14 The DDE for each FDA-approved drug associated with pancreatic cancer and the literature mining results**

This table lists all the DDE scores calculated using NOGEA. The result of literature mining contains the number of articles derived by searching each drug name, “pancreatic cancer” as well as both search terms, respectively. The P-values were assessed using the hypergeometric test.

**Table S15 Differentially expressed genes and the predicted effected genes after treatment with Saquinavir and Celecoxib.**

CELDEG: the differentially expressed gene after treatment with Celecoxib. CELPEG: the predicted effected gene after treatment with Celecoxib. SAQDEG: the differentially expressed gene after treat with Saquinavir. SAQPEG: the predicted effected gene after treatment with Saquinavir.

**Table S16 Release versions of the database used in this study.**

This table lists all the databases and corresponding versions that were used in this study.

